# Protein-S-Nitrosylation of Human Cytomegalovirus pp65 Reduces its Ability to Undermine cGAS

**DOI:** 10.1101/2024.11.20.624495

**Authors:** Justin B. Cox, Masatoshi Nukui, Eain A. Murphy

**Affiliations:** Microbiology and Immunology Department, SUNY-Upstate Medical University, Syracuse NY 13210

## Abstract

Post-translational modifications (PTMs) are key regulators of various processes important for cell survival. These modifications are critical for dealing with stress conditions such as those observed in disease states and during infections with various pathogens. We previously reported that during infection of primary dermal fibroblasts, multiple Human Cytomegalovirus (HCMV) encoded proteins were post-translationally modified by the addition of a nitric oxide group to cysteine residues, a modification called protein-S-nitrosylation. For example, tegument protein pp71 is nitrosylated, diminishing its ability to inhibit STING, a protein necessary for DNA virus immune response. Herin, we report that an additional HCMV tegument protein, pp65, responsible for the inhibition of cGAS, is also modified by protein-S-nitrosylation on two cysteine residues. Utilizing site-directed mutagenesis to generate recombinant viruses that encode a pp65 that cannot be protein-S-nitrosylated, we evaluated the impact of this PTM on viral replication and how the virus impacts the cGAS/STING pathway. We report that the nitrosylation of pp65 negatively impacts its ability to block cGAS enzymatic functions. pp65 protein-S-nitrosylation mutants demonstrated a decrease in cGAS/STING induced IRF3 and TBK1 phosphorylation. Additionally, we observe a reduction in IFN-β1 secretion in NuFF-1 cells expressing a nitrosylation-resistant pp65. We report that HCMV expressing a protein-S-nitrosylation deficient pp65 is resistant to the activation of cGAS in the infection of primary dermal fibroblasts. Our work suggests that nitrosylation of viral proteins may serve as a broadly neutralizing mechanism in HCMV infection.

**Importance:** Post translational modifications (PTM) are utilized by host cells to limit an invading pathogen’s ability to establish a productive infection. A potent PTM called protein-S-nitrosylation has anti-bacterial and anti-viral properties. Increasing protein-S-nitrosylation with the addition of nitric oxide donor compounds, reduced HCMV replication in fibroblasts and epithelial cells ^1^. We previously reported that protein-S-nitrosylation of HCMV pp71 limits its ability to inhibit STING. Herein, we report that the protein-S-nitrosylation of HCMV pp65 impacts it’s ability to limit cGAS activity, an additional protein important in regulating interferon response. Therapeutically, patients provided nitric oxide by inhalation reduced viral replication in COVID-19, influenza and even impacted bacterial growth within patients lungs ^2,3^. It is thought an increase in free nitric oxide increases the frequency of nitrosylated proteins ^4^. Understanding how protein-S-nitrosylation regulates a common DNA virus like HCMV will provide insights into the development of broadly neautralizing therapeutics in drug resistant viral infections.

## Introduction

Human Cytomegalovirus (HCMV), a large ubiquitous beta-herpesvirus, is endemic in the human population ^5,6^. Upon infection with HCMV, the virus persists for the lifetime of the host as the virus has the capacity to establish a latent infection within myeloid progenitor cells coupled with sporadic reactivation resulting in lytic replication in various cell types ^7,8^. It is estimated that 30-80% of adults are HCMV seropositive and, typically, a competent immune system renders lytic reactivations asymptomatic ^9,10^. However, in immunocompromised individuals, such as bone marrow recipients and HIV-positive patients, HCMV often results in severe disease and death ^11,12^. Equally problematic are HCMV infections in neonates with underdeveloped immune systems. HCMV is currently the leading cause of congenital infections and is responsible for cognitive and physical birth defects such as hydrocephaly and microcephaly in newborns if the mother contracts a primary HCMV infection while pregnant ^13^. Currently, there is no vaccine for HCMV, thus making therapeutic interventions for pre-existing infections a focus for limiting infection spread. While direct acting antiviral drugs have proven effectual for controlling HCMV infection, they are becoming less clinically effective as resistant strains of the virus have emerged ^14,15^. Thus, identifying how the innate and intrinsic immune responses limit HCMV infection and characterization of PTMs that undermine viral virulence factors is essential in developing novel neutralizing therapeutics.

Innate immunity is one of the earliest steps in combating viral and bacterial infections. Pathways involved in this recognition are responsible for identifying pathogen-associated molecular patterns (PAMPs) within the cytoplasm of a cell. The recognition of PAMPs, such as cytoplasmic dsDNA, by pattern recognition receptors (PRRs) like the cyclic guanine-adenosine synthase (cGAS) enzyme is essential for controlling DNA virus infections ^16–19^. Once cGAS encounters cytoplasmic dsDNA, it facilitates the enzymatic reaction fusing ATP and GTP thereby generating guanosine-adenosine 2’3’-cyclic monophosphate (2’3’-cGAMP), an agonist for <u>st</u>imulator of <u>in</u>terferon <u>g</u>enes (STING) ^16,20–25^. Stimulation of cGAS results in the potent inhibition of viruses such as HCMV ^26–28^. However, HCMV encodes various proteins that function to undermine cellular DNA sensors such as STING and cGAS ^10,29^. Packaged in the tegument of HCMV is pp65, a protein encoded by open reading frame Unique Long 83 (UL83) ^30^. The direct interaction of pp65 and cGAS decreases the latter’s ability to produce 2’3’-cGAMP thus reducing STING-mediated immunity and interferon response 30,31. Underscoring its importance during the initial stages of viral infection, pp65 comprises approximately fifteen percent of the tegument of infectious virions and most of the approximate weight of dense bodies ^32^.

Intrinsic immunity not only detects and responds to foreign substances, it also functions to induce modifications viral proteins leading to a reduction in the protein’s activity. A cell typically modifies the biological functions of proteins through PTMs, such as modifications that may lead to protein degradation, thus limiting their presence within an infected cell. Modifications such as ubiquitination, SUMOylation, and phosphorylation can each serve as innate immune mechanism for infected cells for virally infected cells 33–37. One such modification that remains understudied due to its liable nature is called protein-S-nitrosylation. This modification involves a nitric oxide group being attached to the thiol group within a cystine amino acid ^38^. This modification can result in multiple consequences such as controlling di-sulfide linkages, altering enzymatic activity, and functioning in anti-viral properties ^38–43^. Zell *et al.* observed that the active sites of the 2A and 3C proteases of coxsackie B3 virus are protein-S-nitrosylated and as a result, this PTM impacts their proteolytic functions ^43^. The anti-viral functions of protein-S-nitrosylation of viral proteins are a novel cellular mechanism that cells can use to limit the functions of the target protein.

We previously reported that protein-S-nitrosylation of HCMV pp71 inhibits its ability to attenuate STING. As we observed that pp65 also is modified in a similar fashion coupled with the fact that it is involved in the cGAS/STING pathway, we wanted to investigate the impact of protein-S-nitrosylation on this protein. Herein, we report that pp65 constructs, in which the identified protein-S-nitrosylated cystines were changed to structurally related serines, inhibit cGAS with better efficiency suggesting that the PTM is inhibitoy to the viral proteins fuctions. Further, we observed that a double mutant pp65 (DM-pp65), that is not protein-S-nitrosylated has a growth advantage over wild-type (WT) pp65 in the presence of a potent cGAS agonist, G3-YSD. This data suggests that protein-S-nitrosylation is not just a singular anti-viral mechanism, but functions as a broadly neutralizing mechanism to improve the function of the cGAS/STING pathway.

## Results

### Human Cytomegalovirus pp65 is nitrosylated in HCMV infection

Our previous work utilized MS/MS analysis to identify peptides that contain a protein-S-nitrosylated cystine during HCMV infection of human Newborn Foreskin Fibroblasts (NuFF-1) and identified 13 HCMV proteins that contained this PTM ^44^. One of these identified proteins was pp65, a tegument protein critical for undermining host innate immune responses. Our analysis results indicated that pp65 is nitrosylated at two distinct cystines sites during infection of NuFF-1 cells (Fig 1A). To validate the protein-S-nitrosylation status of pp65 and characterize the kinetics of when the PTM is added to pp65, human NuFF-1 cells were infected with HCMV at an MOI of 1 and cell lysates were collected in non-reducing conditions to preserve the protein-S-nitrosylation modification at 24, 48 and 72hpi. Equal amounts of lysate were used in a Biotin Switch Assay in which all nitrosylation groups are converted to biotins thereby allowing for modification detection. Peptides containing biotin were affinity purified followed by immunoblot analysis with anti-pp65 and anti-actin antibodies. Actin was specifically chosen as a suitable control as it has been reported to be protein-S-nitrosylated ^45–48^. Our results show that as early as 48 hours post infection (hpi) pp65 is nitrosylated with increasing levels of the PTM being observed at 72 hours hpi (Fig 1B) validating the results of the MS/MS analysis that pp65 is nitrosylated during the course of a lytic HCMV infection and the modification is observed with similar kinetics and levels as the protein expression of pp65.

**FIG 1.**
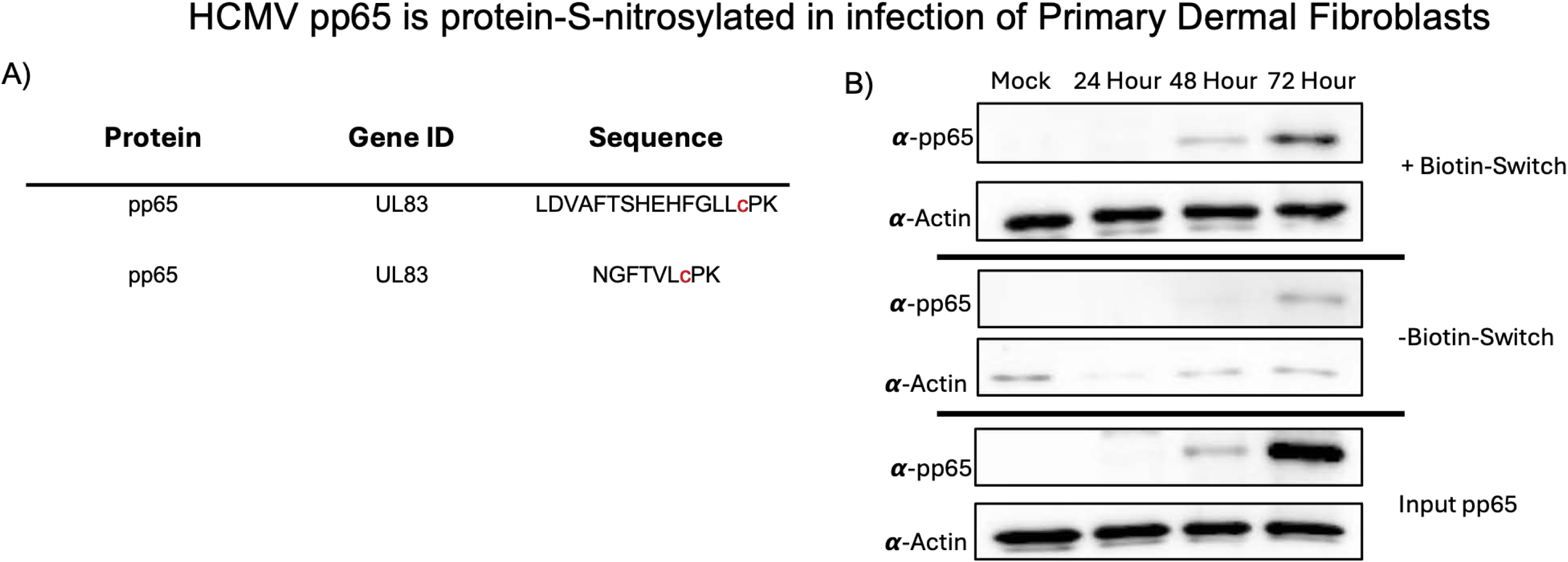
HCMV pp65 is protein-S-nitrosylated in infection of Primary Dermal Fibroblasts. A) Primary Foreskin Fibroblasts were infected with TB40/E-mCherry-UL99-eGFP at an MOI of 1 for 96 hours. After 96 hours cell lysates were collected and protein was subjected to biotin-switch to label nitrosylated proteins with biotin then affinity purified with streptavidin beads. Affinity purified protein was analyzed by unbiased MS/MS analysis identifying 13 viral proteins to be protein-S-nitrosylated. The small red C indicates where a nitrosylation group was detected in our MS/MS analysis. The nistrosylated sites for pp65 only are shown in this figure. B) NuFF-1 cells were infected at an MOI of 1 for 24-,48-, and 72-hours, and 100 ug of protein was subjected to a biotin switch and then affinity purified with streptavidin beads. Biotinylated protein was separated on an 8% SDS-PAGE and transferred to a nitrocellulose membrane blocked in 5% BSA and probed for pp65. A,B) n=3.

### Nitrosylation deficient pp65 expressing HCMV replicates with similar kinetics compared to WT virus

To determine if protein-S-nitrosylation impacts the biological functions of the viral protein, we generated a nitrosylation deficient pp65, termed DM-pp65, where each of the two identified protein-S-nitrosylated cystines were mutated to the structurally similar amino acid serine which is identical to cystine except for the absence of a sulpher atom thus rendering it a non-suitable substrate for the modification. As recombineering may cause spurious off site mutations to arise thereby confounding potential results, we generated a revertant of our mutant DM-pp65 virus so that the two mutated serines we reverted back to the original cystines, termed Rev-pp65. To determine if DM-pp65 exhibits an altered growth kinetics when compared to WT or Rev-pp65 virus, NuFF-1 cells were infected at an MOI of 1.0, and cell-free virus was collected on days 0, 3, 5, and 7 days post-infection (dpi). A fraction of inoculum was saved to confirm the initial starting titer for infection. Viral titers were then quantified by TCid50 analysis. Viral titers of WT, DM-pp65 and Rev-pp65 virus exhibited similar growth kinetics suggesting that the mutations had little impact on the ability of the virus to undergo lytic replication in fibroblasts (Fig 2A).

**FIG 2.**
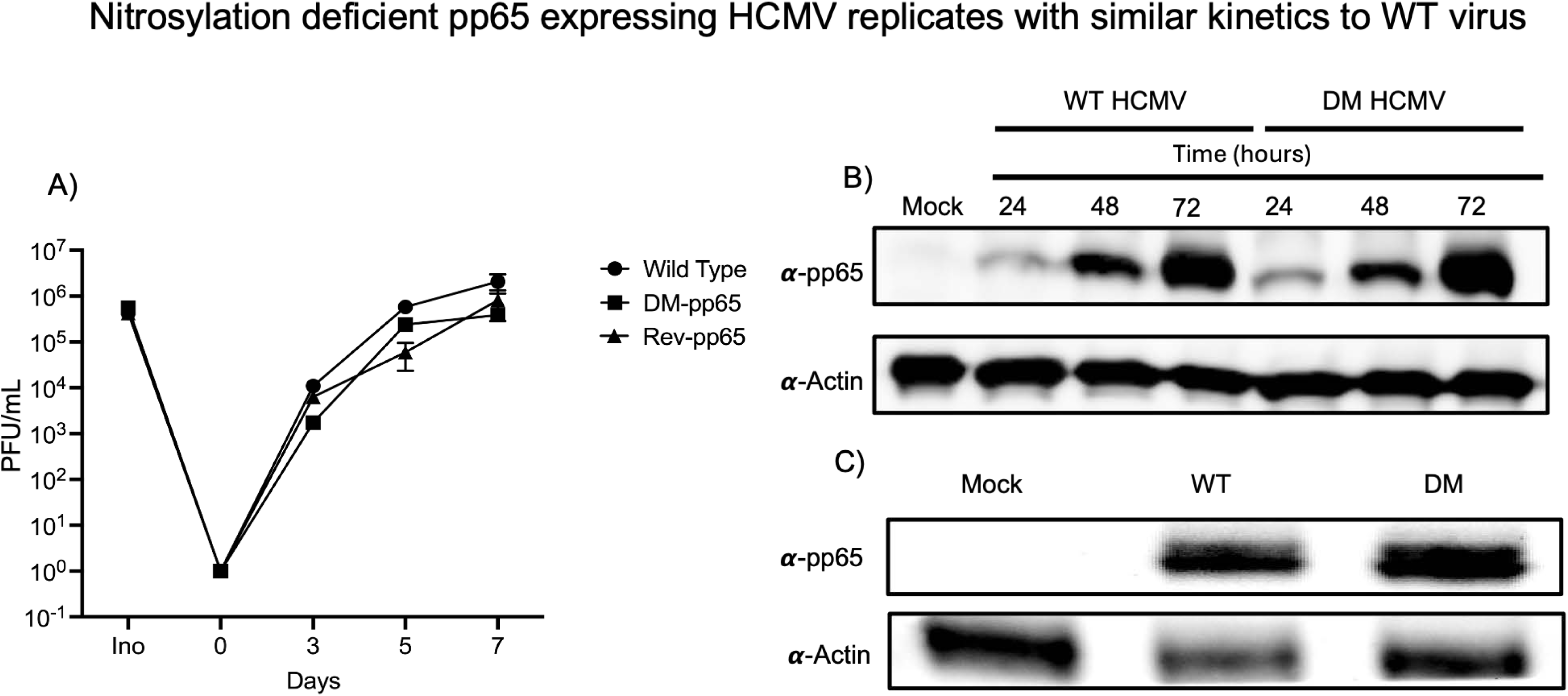
Nitrosylation deficient pp65 expressing HCMV replicates with similar kinetics to WT vims. A) HFF-1 cells were infected at an MOI of 1 with WT, DM, or Revertant HCMV and supernatant was collected days 0-,3-,5-, and 7 dpi. Viral titers were measured by TCID50 with mCherry expression. Inoculum was saved to determine equal PFU upon infection of NuFF-1 cells. B) NuFF-1 cells were infected with WT or DM pp65 at an MOI of 1 and cell lysates were collected at 0-,24-,48-, and 72 hpi. 30 ug of protein was separated on and 8% **SDS-PAGE,** transferred to a nitrocellulose membrane and blocked with 5% BSA. The membrane was then probed with anti-pp65 and anti-actin. C) HFF-1 cells were infected at an MOI of 1 with WT or DM-pp65 HCMV for 6 hours. 30 ug of protein was separated on an 8% **SDS-PAGE,** transferred to a nitrocellulose membrane and blocked with 5% BSA. Membrane was then probed for anti-pp65 and actin. A,B,C) n=3.

Next, we wanted to evaluate the protein kinetics of pp65 expression in WT and DM-pp65. pp65 is a viral late transcript with observable *de novo* expression initiating at 24 hours post infection and highest expression observed late in infection ^49–51^. To compare the *de novo* expression of WT and DM-pp65 NuFF-1 cells were infected at an MOI of 1 and cell lysates were collected at 24, 48, and 72 HPI. Protein expression was then measured by immunoblot. At each time point, we observed similar levels of pp65 protein expression from the WT and DM-pp65 viruses, suggesting that the mutations did not impact the levels of pp65 protein expression or the timing of their translation (Fig 2B).

pp65 is a major tegument protein important in the initial stages of infection and represents up to 15% of the mass of an infectious virion ^52^. To determine if protein-S-nitrosylation impacts the loading of pp65 into the tegument of HCMV, NuFF-1 cells were infected at an MOI of 1.0 with WT and DM-pp65 and lysates collected at 6hpi, a timepoint prior to the *de novo* synthesis of pp65. Lysates were then collected, and deposition of tegument delivered pp65 were analyzed by immunoblot. We observed similar band intensities in both WT and DM-pp65 conditions suggesting that loading and/or delivery of pp65 is not impacted by the substitutions of serine for cystines in pp65 (Fig 2C). In sum, the substitution of serine for protein-S-nitrosylated cystine did not alter the replication kinetics or tegument loading of pp65 in fibroblasts.

### DM-pp65 HCMV replication is more resistant to G3-YSD treatment

A key biological function of pp65 is to bind to and inhibit the activity of cGAS ^30^. We, therefore, focused our efforts on evaluating the impact of protein-S-nitrosylation of pp65 in the context of the cGAS/STING pathway, as we have previously reported that protein-S-nitrosylation of another HCMV tegument protein, pp71, reduced its ability to inhibit the biological functions of the STING protein. cGAS upon engagement with cytoplasmic dsDNA produces 2’3’-cGAMP which in turn stimulates STING to initiate interferon and pro-inflammatory cytokine transcriptional induction. To activate cGAS, we treated cells a cGAS agonist, G3-YSD, that was introduced into cells by lipid based transfection. We monitored cell viability upon treatment of G3-YSD using a WST viability assay. Cell viability remained consistent with no significant reduction of viability observed from 0.4 to 0.8ug of G3-YSD no toxicity was observed. No significant toxicity was observed in the vehicle group which was just lipofectamine. However, there was a significant reduction in cell viability at 1.6ug of G3-YSD with a cell viability of approximately 83% (Fig 3A). Using a cGAS agonist, G3-YSD, we observed a dose dependent inhibition of HCMV virion production following treatment with G3-YSD at concentrations that did not alter cellular viability highlighting the role of activated STING on establishing an antiviral state (Fig 3B). Since decreases in viral titers were identified in WT HCMV infection, we wanted to investigate how protein-S-nitrosylation deficient HCMV DM-pp65 replicates in the presence of an activated anti-viral state. NuFF-1 cells were pre-transfected with 0.4,0.8 and 1.6 ug of G3-YSD or just vehicle as 1.6 ug of G3-YSD was sufficient in reducing WT HCMV viral titers below the limit of detection. NuFF-1 cells were then infected at an MOI of 0.1 with WT or DM-pp65 HCMV and media collected 7 dpi and titers measured by TCid50 assay. As expected, percent viral titer decreased in WT HCMV infection at all concentrations of G3-YSD as seen in figure 3B. In the DM-pp65 infected cells there was no significant decrease in viral titers treated with 0.4, and 0.8 ug of G3-YSD (Fig 3C). However, a significant decrease in percent viral titers in the 1.6 ug treatment group did attenuate DM-pp65 replication by 50% but not below the limit of detection like WT HCMV. This data suggests that protein-S-nitrosylation deficient DM-pp65 can antagonize cGAS even in an activated state whereas WT HCMV infection is inhibited. This result would lead one to hypothesize that DM-pp65 may be degrading cGAS or that the protein-S-nitrosylation of pp65 may cause a reduced interaction causing a stronger interaction between the cGAS and DM-pp65.

**FIG 3.**
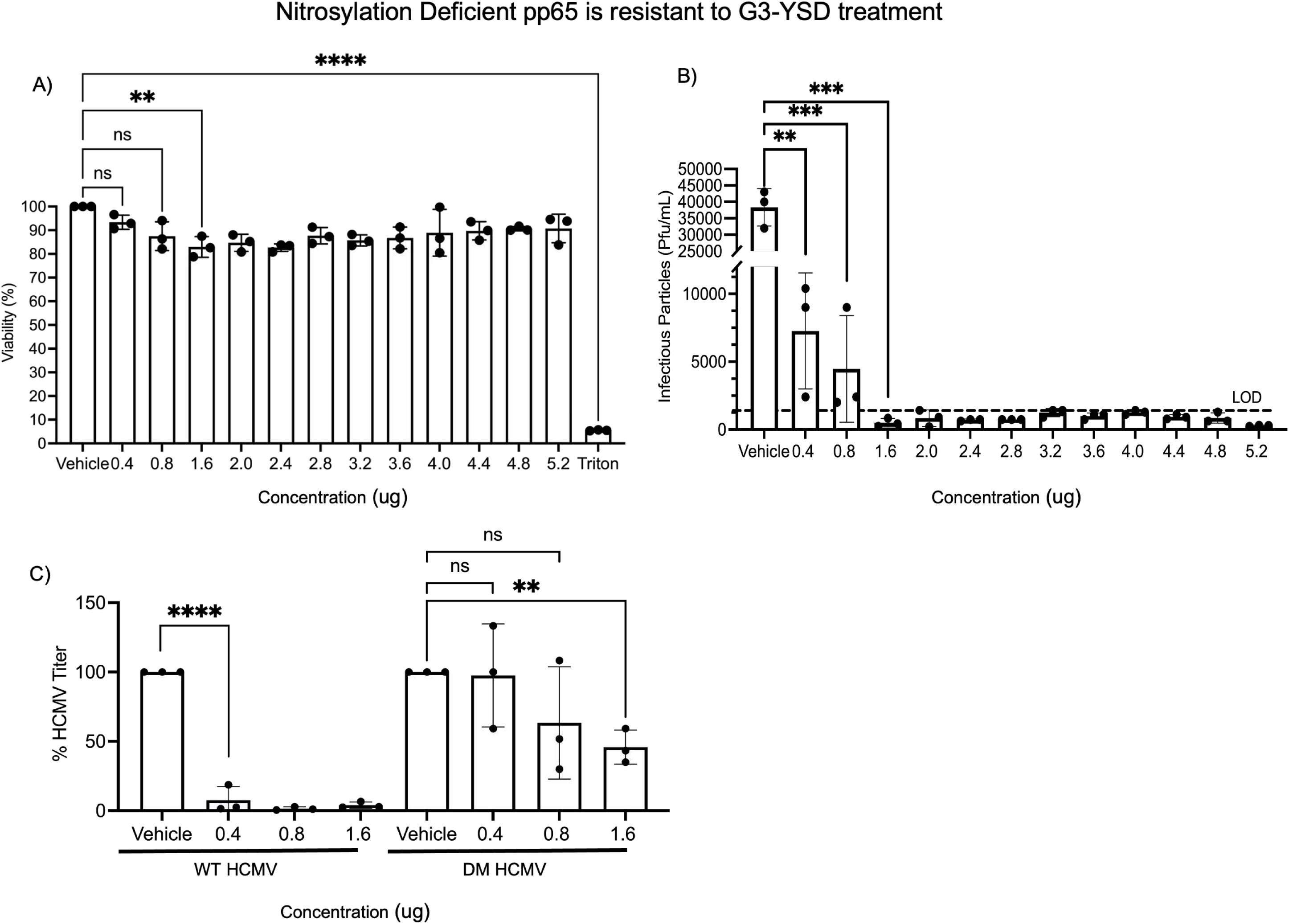
Nitiosylation Deficient pp65 is resistant to G3-YSD treatment: A) WST assay of NuFF-1 cells transfected with G3-YSD 24 hours after transfection. NuFF-1 cells were transfected for 6 hours, media removed and 24 hours later subjected to WST assay. 10% triton X-100 was used as a control to observe cell death within the assay. B) NuFF-1 cells were transfected with G3-YSD for 6 hours with indicated concentrations and then infected with WT HCMV 24 hours later at an MOI of 0.1. Viral titers were measures by TCID50 assay 7 days post infection and reported as PFU/mL. C) NuFF-1 cells were transfected with 0.4, 0.8, and 1.6 ug of G3-YSD for 6 hours and then 24 hours later infected with WT or DM-pp65. All viral titers were measured 7 days post infection by TCID50 assay. A,B,C) n=3, p<0.01*“,p<0.001*”“, p<0.0001”*“.

### cGAS is not degraded during infection of NuFF-1 cells with either WT or DM-pp65

DM-pp65 demonstrates increased resistance to cGAS activation as notated from its ability to replicate in the presence of G3-YSD with a possible explanation that the protein levels of cGAS are regulated by pp65. To determine if cGAS is degraded in WT or DM-pp65 HCMV infection, NuFF-1 cells were infected at an MOI of 1 with WT and DM-pp65 and lysates were collected at 12, 24, 48, and 72 HPI. Immunoblots using cGAS specific antibodies indicated that in both the WT and DM-pp65 infected NuFF-1 cells degradation of cGAS was not observed at any of the time-points of infection (Fig 4). This was in line with previous reports that cGAS is not degraded during HCMV infections where wildtype pp65 is expressed ^30^. This data suggests that the resistance observed in DM-pp65 HCMV is not from the degradation of cGAS but possibly limitation of the activity of cGAS with better efficiency then a WT pp65.

**FIG 4.**
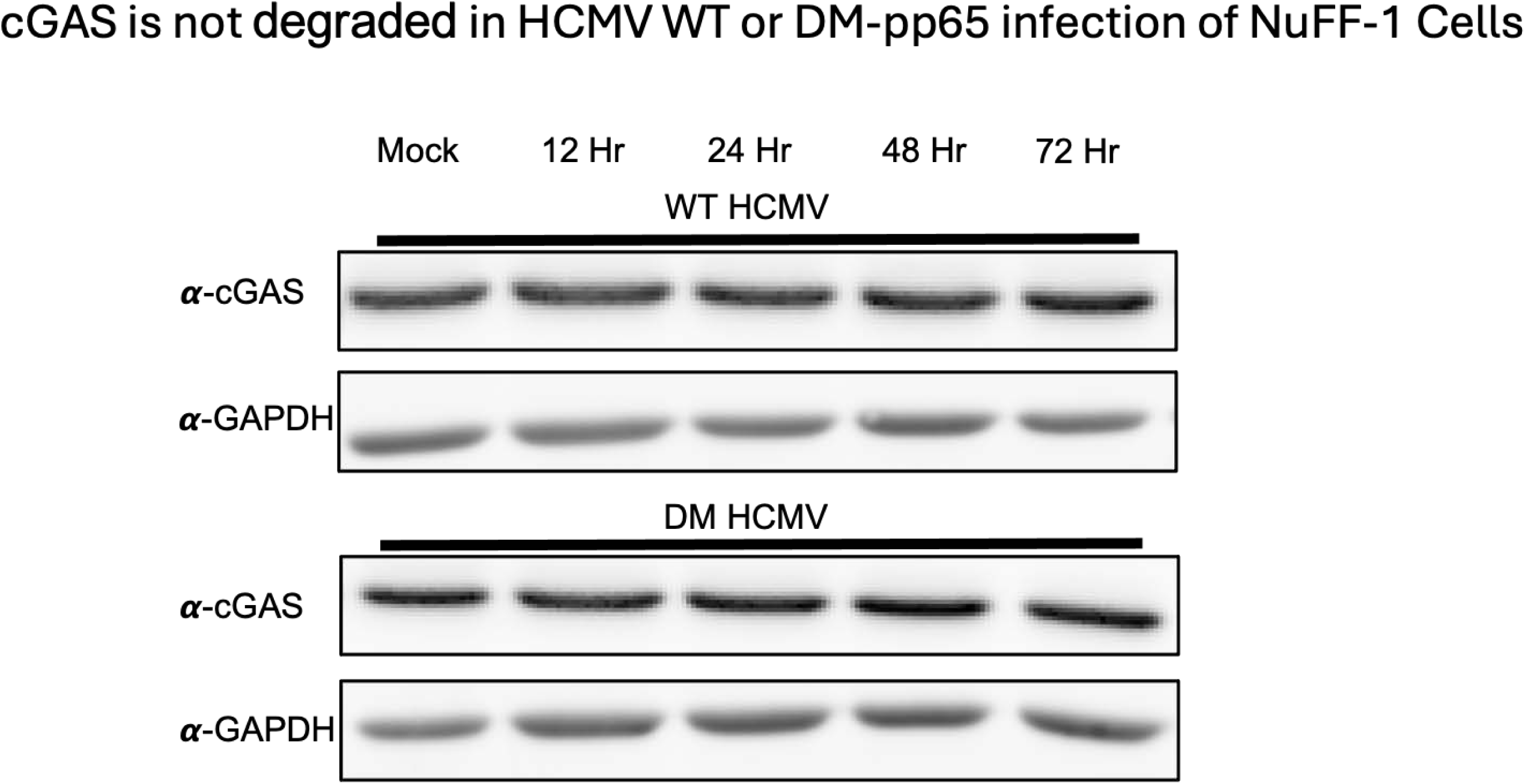
cGAS is Not Degraded in HCMV WT or DM-pp65 Infection of Pflmary Dermal Fibroblasts: NuFF-1 cells were infected with HCMV WT or DM-pp65 at an MOI of 1 and cell lysates were collected at 12-, 24-, 48- and 72 hpi. 30 ug of protein was separated on ari 8°/oSDS-PAGE and transferred to a nitrocellulose membrane. Blots were probed for anti-pp65 and GAPDH as a loading control. Blots were stained with secondary antibody and then imaged with HRP and GAPDH was imaged in a rhodamine channel. n=3.

### Wild-Type and DM-pp65 interact with cGAS with similar efficiencies

The data thus far supports a model in which the nitrosylation of pp65 may serve as an anti-viral property due to its reduced capacity to block the activation of the STING pathway in the presence of G3-YSD. However, it is critical to understand the biological impact of pp65 when protein-S-nitrosylation of the protein is inhibited. To test this we generated stable cell lines that express three nitrosylation resistant pp65 mutants SM1- pp65, SM2-pp65 and DM-pp65. We only determined if WT-pp65 stable cell lines are protein-S-nitrosylated by biotin-switch assay as SM1 and SM2 still have cystines that can be protein-S-nitrosylated. As expected, immunoblotting identified a band in the avidin purified group that underwent the biotin switch. This suggests that pp65 is still protein-S-nitrosylated in stable cell lines without the presence of other viral factors and viral DNA (Fig 5A).

**FIG 5.**
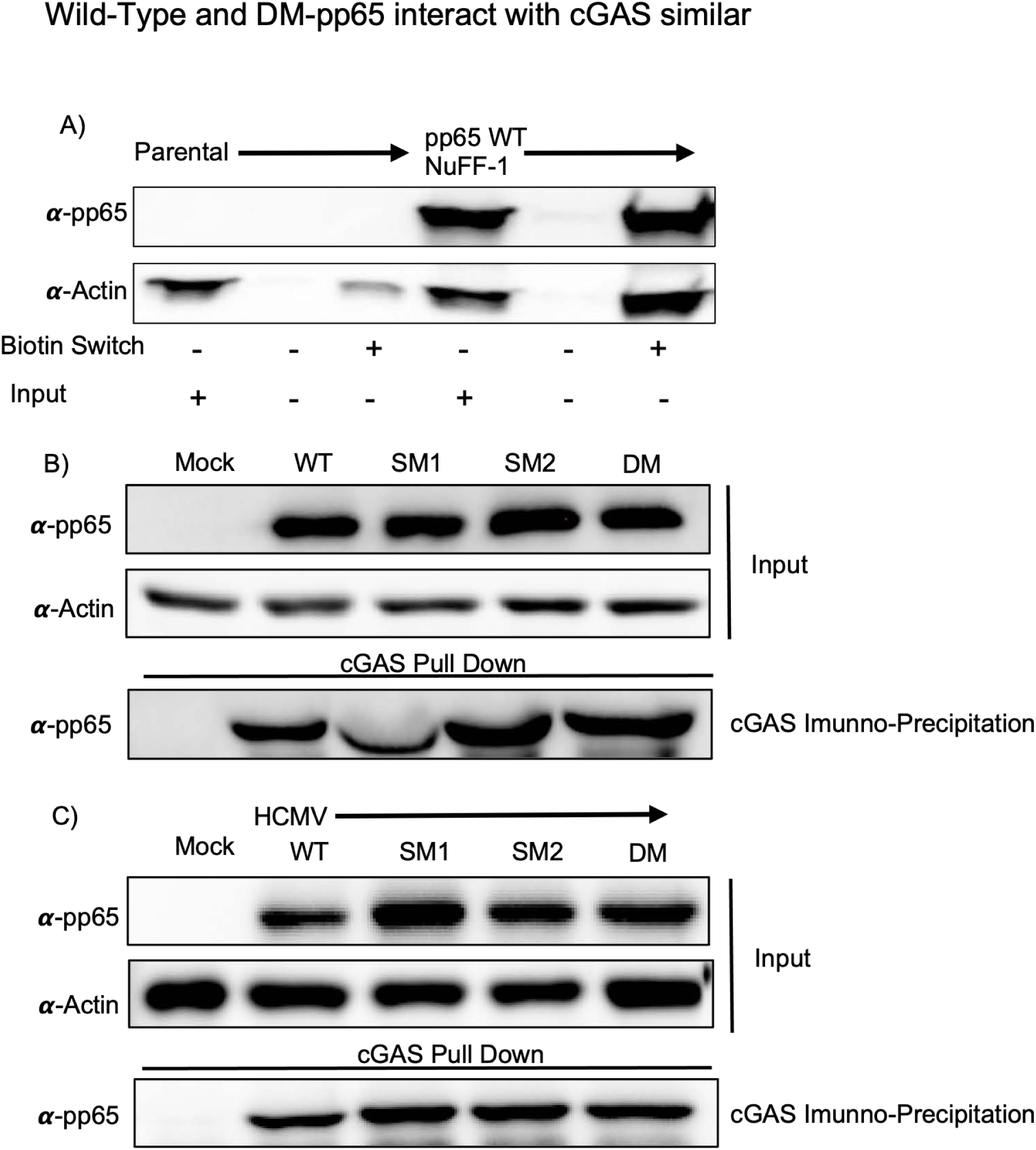
HCMV pp65 mutants interact with cGAS in stable cell lines and infection: A) NuFF-1 parental and pp65 WT expressing NuFF-1 cells were lysed using the protein-S-nitrosylation detection kit. Proceeding cell lysis, 100ug of cell lysate underwent the “biotin-switch” reaction following the manufacturer’s instructions. Following the biotin switch all reactions were affinity purified with avidin-coated beads. Protein was then separated to an 8% **SDS-PAGE** and blotted with primary antibody specific to pp65. B) Stable cell lines were plated at 85% confluency and the next day lysed with Co-IP lysis buffer. Protein was incubated with cGAS coupled dynabeads overnight at 4 degrees Celsius. Protein was separated by an 8% SDS-PAGE and 5 ug of input lysate was used to confirm expression of pp65 protein. The membrane was then probed with pp65 antibody to confirm pulldown from interacting with cGAS. Actin was used to confirm equal loading of input. C) HFF-1 cells were infected at an MOI of 3 for 72 hours with WT and all pp65 nitrosylation deficient mutants. Cells were lysed with Co-IP lysis buffer and protein was incubated with cGAS coupled dynabeads overnight at 4 degrees Celsius. 5 ug of input was separated as well to confirm equal expression of pp65 in each reaction. Protein was separated on an 8% SDS-PAGE and protein was transferred to a nitrocellulose membrane. The membrane was then probed with pp65 antibody to confirm pulldown from interacting with cGAS. Actin was used to confirm equal loading of input. A,B,C) n=3.

We now wanted to determine if additional viral proteins are required for the interaction of pp65 and cGAS by immunoprecipitating cGAS with WT-pp65 and all nitrosylation-deficient mutants. To determine if wildtype and mutant pp65 proteins can still interact with host encoded cGAS, NuFF-1 pp65 expressing cells were lysed with Co-IP lysis buffer and the immunoprecipitation was repeated as above. In the absence of other viral factors, we observed robust interaction with cGAS and wildtype pp65 as well as each of the protein-S-nitrosylation deficient mutants (Fig 5B). This data suggests that blocking protein-S-nitrosylation of pp65 has little impact on its interaction with cGAS and the ability of the DM-pp65 virus to show increased replication in the presence of activated cGAS is likely due to a different effect than interaction with cGAS.

HCMV pp65 interacts with cGAS thereby inhibiting it’s capability to produce 2’3’-cGAMP reducing STING activation ^30^. Defining if this interaction occurs when the nitrosylation of pp65 is blocked would explain the increased replication of DM-pp65 in the presence of G3-YSD. NuFF-1 cells were infected with WT, SM1-pp65, SM2-pp65 or DM-pp65 expressing HCMV at an MOI of 1 for 72 hours. SM1-pp65 and SM2-pp65 are viral mutants in which only one of the two identified protein-S-nitrosylated cystines were mutated to a serine so these viruses can still be protein-S-nitrosylated at one site. Cells were lysed at 72 hours and 50 ug of protein was incubated with anti-cGAS coupled beads. Immunoblotting of the IP revealed bands of equal protein densities for all pp65 mutants (Fig 5C) suggesting that the protein-S-nitrosylation status of pp65 did not alter the ability of the viral protein to interact with cGAS.

### DM-pp65 antagonizes cGAS with better efficiency than wild type pp65 in stable cell lines

To determine if DM-pp65 inhibits cGAS activation to a different level than that of the wildtype protein, stable cell lines expressing either WT or DM-pp65 were transfected with the cGAS activator, G3-YSD, and cell lysates were collected at 2 and 4 hours post transfection. As expected, there was an increase in the levels of phosphorylation of IRF3 and TBK1 in the parental cell lines and a decrease in the phosphorylation status of both proteins in the presence of WT pp65. Importantly, in the DM-pp65 expressing NuFF-1 cells there was a reduction of phosphorylated IRF3 and TBK1 levels when compared to WT expressing pp65 NuFF-1 cells (Fig 6A,B). This decrease of phosphorylation suggests that this would result in a decrease in downstream antiviral factors such as the secretion of IFN-β1. To determine if there are changes in IFN-β1 secretion, stable cell lines were transfected with 0.2 ug of G3-YSD and then the supernatant was collected 8 hours post treatment to be analyzed by Enzyme-Linked Immunosorbent Assay (ELISA). The cell line expressing DM-pp65 group showed a significant decrease in the secretion of IFN-β1 post G3-YSD treatment compared to parental or WT pp65 expressing cell lines (Fig 6C). Our data reveals that protein-S-nitrosylation deficient pp65 antagonizes cGAS with better efficiency and leads to a decrease of phosphorylation of IRF3 and TBK1 and a reduction in IFN-β1 secretion. This observation suggests that protein-S-nitrosylation of viral proteins like pp65, coupled with our earlier findings with pp71 suggests that this modification may function as a broadly neutralizing mechanism in HCMV infection.

**FIG 6.**
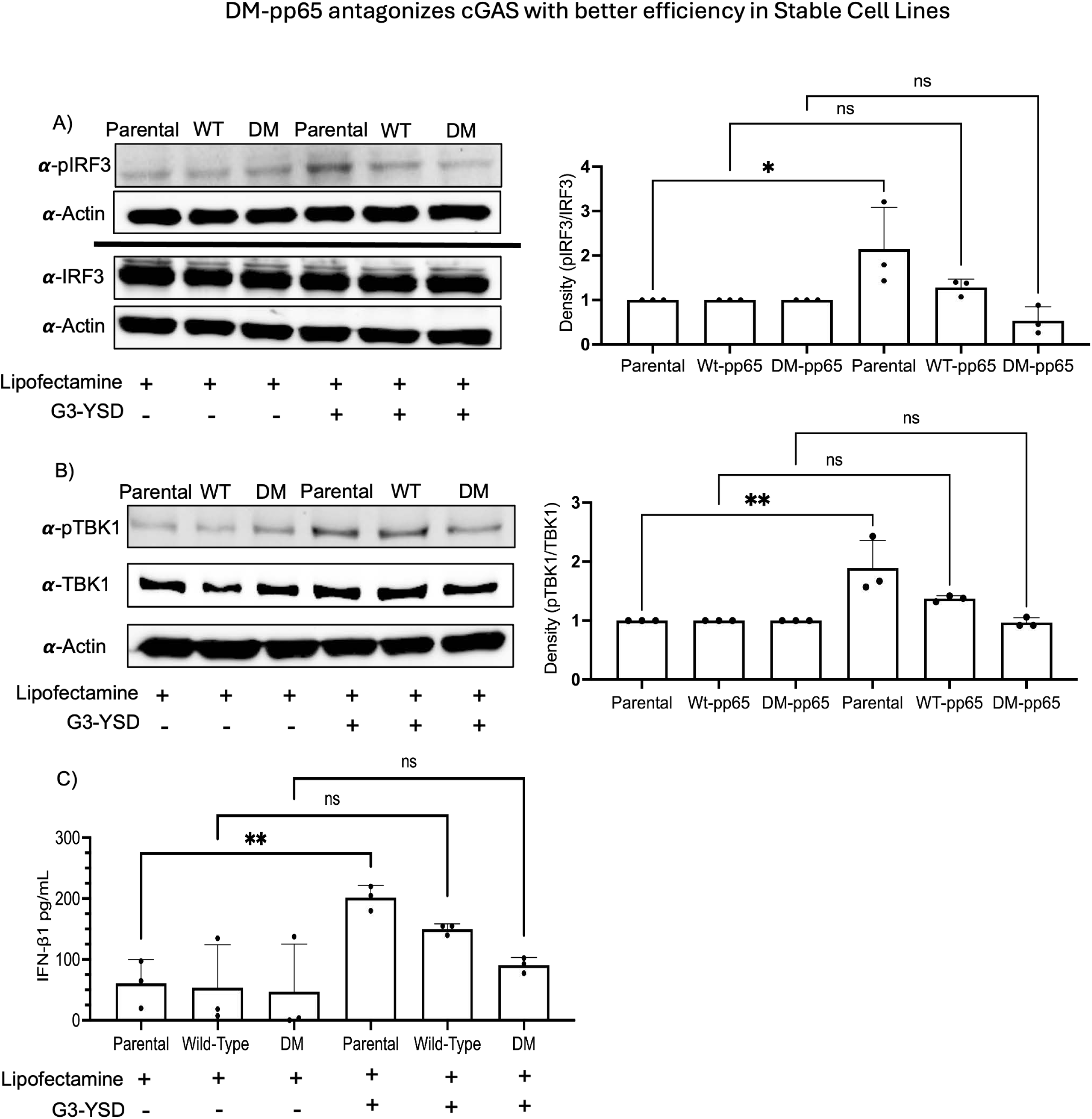
The presence of DM-pp65 in NuFF-1 cells decreases the phosphorylation of plRF3 and TBK1 after G3-YSD treatment: A) NuFF-1 parental and ie cells were transfected with 0.2 ug of G3-YSD or vehicle for 2 hours and then cell lysates were collected, processed and 30 ug of protein was separated on an 8% SDS-PAGE. Blots were probed with anti-TBK1 and pTBK1 and then multiplexed with fluorescent secondary antibodies. B) NuFF-1 and stable cells were transfected with 0.2 ug of G3-YSD for 4 hours and then cell lysates were collected, processed and 30 ug of protein was separated on an 8% **SDS-PAGE.** Blots were probes with anti-lRF3 and pIRF3 and anti-rabbit HRP was used to develop blots to measure the expression of phosphorylation groups. C) NuFF-1 cells parental and stable cell lines were transfected with 0.2 ug of G3-YSD for 8 hours and then supernatant collected. Supernatant was then analyzed for IFN-β1 secretion by ELISA. OD readings were read at 450 nm. A,B,C) n=3, p<0.05*, p<0.01**.

**FIG 7.**
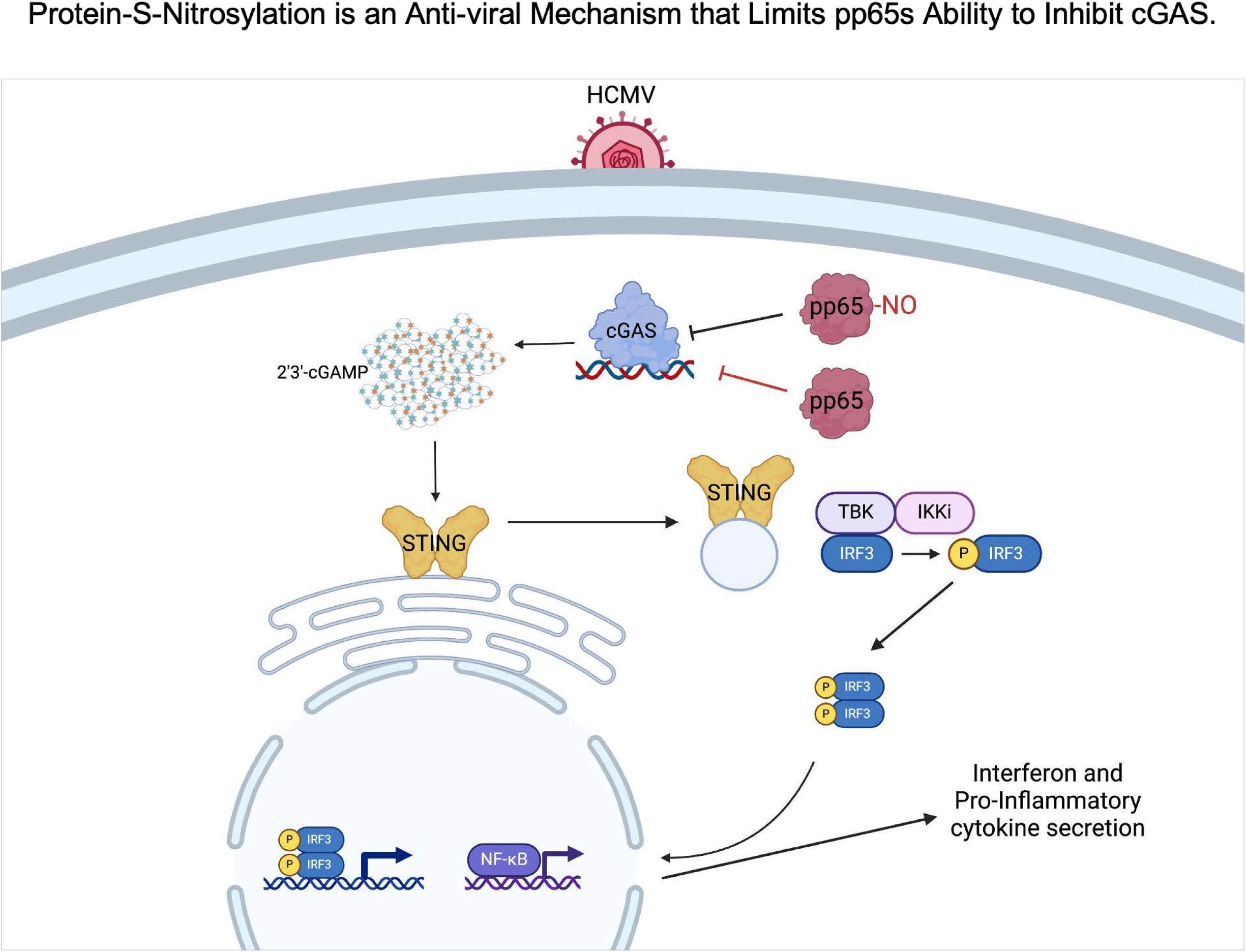
The nitrosylation of pp65 is an antiviral mechanism limiting its ability to inhibit cGAS. Our data suggests that when nitrosylation of pp65 is blocked it has a better ability to antagonize cGAS. We recognized this by a decrease of phosphorylation of IRF3 and TBK1 in the presence of the DM-pp65 mutant protein, this greater decrease of IRF3 phosphorylation led to a decrease in IFN-β1 secretion as well. We observed DM-pp65 is resistant to cGAS activation in NuFF-1 cells, having a fitness compared to WT virus. Our model introduces the idea that nitrosylation of viral proteins being an anti-viral mechanism is not specific to pp71 but also limits the ability of pp65 suggesting that nitrosylation is a potent and broadly neutralizing mechanism in HCMV.

## Discussion

Using mass spectrometry based proteomics, we identified 13 viral proteins in HCMV infection to be protein-S-nitrosylated. We reported that when protein-S-nitrosylation of pp71 is inhibited it results in an increased antagonization of the STING pathway compared to WT pp71, thus suggesting the PTM blocks functions of the viral protein. We wanted to understand if this PTM may serve as a universal anti-viral mechanism and not just the inhibition of one specific viral protein. Herein, we focused on identifying the impact of protein-S-nitrosylation of pp65 as our MS/MS analysis identified this critical HCMV protein to be nitrosylated at two independent cystines and pp65 inhibts the cytosolic DNA sensor cGAS, a key activator of STING. We observed that by 48 hpi pp65 is protein-S-nitrosylated. Robust protein expression of pp65 begins at 24 hpi but we did not observe significant levels of protein-S-nitrosylation of pp65 accumulating at this earlier time point (Fig 1B) suggesting that this modification may be an induced response to viral infection.

Site-directed mutagenesis with BAC recombineering is an important technique allowing one to change specific amino acids in a protein in order to study specific phenotypes. Two sites identified to be protein-S-nitrosylated within pp65 were changed from cystines to the structurally similar serine in order to understand the biological impact where pp65 protein-S-nitrosylation is blocked. Often mutating a critical protein may cause a growth defect in viral replication as it may impact key functions of the protein. A single-step growth curve comparing the replication of WT, DM-pp65 and a revertant virus in which the mutation was reverted back to the wildtype cystines did not reveal a growth disadvantage for the DM-pp65 virus as it grew with similar kinetics as the WT and Rev HCMV suggesting that the induced mutations in pp65 did not impact lytic replication in fibroblasts (Fig. 2A). As mutations can alter the folding and/or turnover of a protein we wanted to evaluate the integrity of pp65 protein expression. No difference was observed in pp65 protein expression between the WT and DM-pp65 virus produced in NuFF-1 cells. However, another important function of pp65 is the loading of viral tegument.

The loading of HCMV proteins like UL25, 41, 43, and 71 are reliant on the expression of pp65 ^52^. The incorporation of proteins into the virion like UL69 and 97 also are impacted by the expression of pp65 ^53^. Since two sites of pp65 were mutated, we next investigated if the loading of the tegument is impacted. NuFF-1 cells infected with WT and DM-pp65 at an MOI of 1 had no difference in the loading of pp65 within the tegument (Fig 2C), yet it remains to be determined if tegument delivered pp65 is nitrosylated, a prospect that is difficult due to protein quantities however. Determining if the protein-S-nitrosylation status of pp65 impacts the loading of other proteins also remains unknown and warrants further investigation, however, as there was no impact on viral lytic replication, this prospect seems unlikely at least for fibroblast infections.

Although the viruses replicate in a similar manner, our focus shifted to investigate if the protein-S-nitrosylation status of pp65 alters its biological functions in attenuating host antiviral responses. We tested the hypothesis that DM-pp65 exhibits a growth advantage in an induced anti-viral state, such as activating cGAS. To this end fibroblasts cells were pre-stimulated with a synthetic double stranded DNA molecule, G3-YSD, that stimulates cGAS activity prior to infection with either WT or DM-pp65 virus. Under these conditions, WT HCMV replication was attenuated by G3-YSD treatment (Fig 3B,C). Importantly, and in support of our model, we observed that DM-pp65 still replicated efficiently even in the presence of an induced anti-viral state (Fig 3C). Although higher levels of G3-YSD treatment did significantly reduce cell viability, tretments with 0.4 and 0.8ug of G3-YSD did not show measurable defects in cell viability and the mutant virus was still resistant in these treatment groups whereas WT HCMV was impeded. It is unlikely that cGAS activation is impacting viral entry into fibroblasts as we still observe mCherry expression in the infected cells as the viruses contain this fluorescent reporter allowing one to monitor infection. Our data suggests that viral exit and egress are impacted as there is a decrease in virion production. While we have not investigated the exact viral lifecycle stage that this inhibition of WT virus occurs it is probable that replication is reduced due to the upregulation of ISGs that inhibit viral replication like IFI16, IFIT1, ISG15, or Mx1 ^54–58^.

A common technique to inhibit the anti-viral state within a cell is for the virus to target immune effectors responsible in limiting viral replication. We wanted to determine if cGAS is degraded in DM-pp65 HCMV infection and if so what mechanisms lead to the degradation of cGAS compared to WT HCMV infection. Interestingly, no degradation of cGAS was observed at any of the time points of HCMV infection in our study (Fig 4). If DM-pp65 degraded cGAS with better efficiency than the WT we would expect to observe a decrease in interferon production from the absence of cellular cGAS and not a reduced activity of cGAS. Concurrent with our results, the Biolatti laboratory which discovered pp65 as a restriction factor for cGAS also observed no degradation of cGAS, and our previous work identified that infection with WT HCMV does not degrade STING^30,44^. This suggests that the pro-viral phenotype we observe when protein-S-nitrosylation is blocked is not due to the degradation of cGAS or STING, but through a different mechanism such as protein-S-nitrosylation impacting pp65’s interaction with cGAS.

To further understand the impact of protein-S-nitrosylation on pp65 in HCMV infection we determined if DM-pp65 demonstrates altered binding to cGAS compared to WT pp65 and if pp65 is protein-S-nitrosylated in the absence of infection. We first observed that in stable cell lines, WT-pp65 is still protein-S-nitrosylated without the presence of other viral factors and without an induced anti-viral state (5A). Biolatti *et al,* reported that pp65 immunoprecipitated with cGAS within protein lysates collected from infection of primary dermal fibroblasts ^30,44^. We observed a similar interaction in our work as well with stable cell lines expressing pp65 and mutant virus (Fig. 5B,C). Interestingly we found that the nitrosylation deficient pp65 mutants also immunoprecipitated with cGAS to similar levels as WT virus infection. A possibility remains that DM-pp65 may lead to a reduction in 2’3’-cGAMP production through reduced enzymatic activity which cannot be measured by Co-IPs and commercial ELISA kits are highly variable in the recognition of 2’3’-cGAMP within cells and remains a challenge. Further, there remains value in testing if the interaction with other cellular innate proteins is impacted from DM-pp65. For example, pp65 inhibits IFI16 and blocks it from bringing DNA to cGAS to produce 2’3’-cGAMP ^59,60^. This could explain why we see no significant alterations in the interaction between cGAS and DM-pp65 but still see a robust replication of virus expressing DM-pp65 after the cGAS/STING pathway is induced.

Next we tested the biological impact of protein-S-nitrosylation of pp65 on downstream antiviral factors of the STING pathway. The stable cell lines were transfected with G3-YSD to induce an anti-viral state inducing canonical cGAS activity. The phosphorylation status of TBK1 and IRF3 were measured and revealed a strong phosphorylation of IRF3 in our parental cells but a decrease of phosphorylation in the infected cells, importantly, with DM-pp65 having a lower level of pIRF3 and pTBK1 when compared to WT (Fig 6A,B). The total protein levels for IRF3, TBK1, and actin were consistently the same indicating that the reduction in phosphorylation was from a mechanism upstream of these factors such as cGAS inhibition. The reduction of TBK1 and IRF3 phosphorylation suggested that DM-pp65 is sufficient to inhibit cGAS, expressing a phenotype where a nitrosylation deficient pp65 has an increased ability to antagonize cGAS. If DM-pp65 influences the phosphorylation of different signaling molecules like NF-κB, which pp65 is known to inhibit, remains unknown and will be a focus of future investigations ^61^.

The phosphorylation of transcription factors like IRF3 leads to the activation of promoters producing cytokines like IFN-β1 and IL-1β. This led us to hypothesize that the reduced level of phosphorylation of IRF3 in the DM-pp65 stable cell lines would lead to a reduction in cytokine secretion. In fact, the DM-pp65 expressing stable cell lines had the lowest amount of IFN-β1 secretion post G3-YSD treatment indicating that protein-S-nitrosylation of pp65 has a biological impact in reducing the activity of pp65 in attenuating the cGAS/STING pathway (Fig 6C). The secretion of IFN-β1 is important in activating dendritic cells which will then activate CD4^+^ T-cells initiating the production of antibodies by B-cells ^62–65^. IFN-β1 can also activate natural killer cells which are important in controlling the reactivation of HCMV ^66–69^. Our results support a model in which protein-S-nitrosylation of viral virulence factors may function is an intrinsic defense mechanism that may assist in more specific antiviral responses like the secondary immune response in viral infections.

Data from this work and from Nukui *et al.* suggests that protein-S-nitroslyation of HCMV viral proteins may serve as a more universal anti-viral mechanism ^44^. The exact factors that promote pp65 and pp71 nitrosylation remains unknown yet nitric oxide, the substrate for protein-S-nitrosylation within HCMV infected cells, has been suggested to have anti-viral properties. When nitric oxide donors like DETA-NONO are provided in trans to HCMV infected cells, viral DNA synthesis is inhibited ^1^. Highlighting the specificity of this response, when HCMV infected cells are treated with both DETA-NONO and nitric oxide scavenger cPTIO viral DNA synthesis returns to normal levels suggesting that the global removal of nitric oxide groups has a pro-viral mechanism ^1^. Nitric oxide donors also reduce the spread of HCMV infection in neural organoids highlighting how potent nitric oxide is to inhibiting HCMV infection in other organs of an infected host ^70^. Nitric oxide did however disrupt the differentiation of neural progenitor cells indicating this treatment may not be appropriate for use in congenital HCMV infections ^70^. We currently do not yet understand why pp65 would conserve this specific amino acid sequence as our data suggests when protein-S-nitrosylation is blocked pp65 antagonizes the cGAS/STING pathway with better efficiency. It may be that a nitrosylated pp65 is important in maintaining latency or is required to interact with other proteins within the cell which we have not investigated yet. These specific cystines may also be important in forming disulfide bridges within the folding of the protein, but we think this is unlikely as the protein still functions with the introduced mutations.

Overall, our work identifies an additional viral protein that is regulated by protein-S-nitrosylation in the cGAS/STING pathway in HCMV infection. We found that blocking protein-S-nitrosylation of pp65 induces a larger decrease in the phosphorylation of IRF3 and TBK1 leading to a decrease in the secretion of IFN-β1. We also observed that DM-pp65 is more resistant to an induced anti-viral state in NuFF-1 cells. Identifying proteins that regulate HCMV infection by protein-S-nitrosylation is important as these factors may be exploited to develop broadly neutralizing anti-viral therapeutics for HCMV infection and potentially other viruses that encode critical proteins which are nitrosylated. Further, utilizing nitrosylation inducing methodologies in a therapeutic manner may provide clinicians with techniques in controlling unmanageable drug resistant viral infections. In sum, our work suggests that protein-S-nitrosylation functions as an antiviral mechanism that could have potentially broad antiviral effects.

## Materials and Methods

### Cell Culture

Newborn Fetal fibroblasts (NuFF-1) and phoenix cells were maintained in Dulbecco’s Modified Essential Medium (Cleveland Clinic) supplemented with 10% Fetal Bovine serum (FBS) (Millipore-Sigma), 1% penicillin-streptomycin solution (Cleveland Clinic) and 2 mM L-Glutamine (Cleveland Clinic). All incubators were maintained at 5% carbon dioxide at 37 degrees Celsius. Cells infected with HCMV for the expansion of virus were supplemented with complete medium that instead of FBS contains 10% newborn calf serum (Gibco). All cells were split with 0.5% Trypsin/EDTA (Cleveland Clinic) at a dilution of 1:2 and fibroblast passages were recorded to limit cellular senescence and limited to no more than 30 passages for experiments.

### Virus propagation

HCMV genomes are contained in bacterial artificial chromosomes (BAC) in SW105 bacteria. For isolation of HCMV BACs, bacteria are streaked on to Luria Broth (LB) Agar-Chloramphenicol plate and grew at 32 degrees Celsius for 24 hours. The bacteria are then cultured overnight in LB-Chloramphenicol broth and the next day BAC DNA is isolated and transfected with electroporation into MRC5 cells that are at a 50% confluency. The next day media is changed on MRC5 cells with complete medium and cultured until the plate is at a 100% cytopathic effect. The virus is then isolated by collecting the supernatant and cells, sonicating, and centrifuging the virus at 20,000 RPM for 1.5 hours at 18 degrees Celsius in a 20 percent sorbitol cushion with an SW-28 rotor in a Beckman-Coulter ultracentrifuge. Virus is then tittered by TCid50 to measure PFU/mL.

### Retroviral Production and Transduction

Phoenix cells were seeded at 80% confluency a day before transfection. To form DNA:Lipofectamine complexes 1ug of retroviral plasmid pLXSN-pp65 WT or DM was incubated with 10 uL of lipofectamine 2000 (Thermo-Fisher) in opti-mem for 30 minutes at room temperature. DNA:Lipofectamine complexes were overlayed onto Phoenix cells overnight and the following day media was changed to 10% NCS supplemented with 2 mM L-Glutamine. The supernatant was collected at 48 and 72 hours post media change, filtered with a 0.45-micron filter (Sigma) and media overlayed on NuFF-1 cells at 70% confluency. NuFF-1 cells were then selected with 300ug/mL of G-418 (Invitrogen) and allowed to grow for two doublings. Protein expression was confirmed with western blot.

### G3-YSD Transfection

G3-YSD was transfected at various concentrations as indicated in the results section. Lipofectamine 2000 (Thermo-Fisher) was incubated with G3-YSD for thirty minutes at room temperature in Opti-Mem to allow complexes to form. After incubation complexes were added to NuFF-1 cells at 75% confluency dropwise and incubated for 2, 4, or 24 hours post transfection depending on the experiment.

### Measuring Viral Titers

Human Cytomegalovirus is tittered by TCid50 assay. NuFF-1 cells are seeded to 90% confluency in a 96-well plate 24 hours prior to infection. Viral supernatant is serially diluted 1:10 and viral plaques counted 14 dpi by visualization of mCherry expression.

### Western Blotting and Co-Immunoprecipitation

Protein was isolated by scraping cells with 100 uL of Pierce RIPA Lysis and Extraction Buffer (25 mM Tris-HCl pH 7.6, 150 mM NaCl, 1% NP-40, 1% sodium deoxycholate, 0.1% SDS; Thermo Fisher Scientific). Lysates were incubated on ice for 30 minutes and sonicated 2X for 15 seconds to lyse cells. Lysates were quantitated by Bradford assay and 30 ug of protein was separated with an 8% SDS-PAGE. Following separation protein was transferred to a nitrocellulose membrane and blocked with 5% BSA for 1 hour at room temperature. The membranes were probed with anti-pp65 (1:200, 8A8), anti-cGAS (1:1000, Sigma), hFAB Rhodamine anti-GAPDH (1:5000, Bio-Rad), anti-actin (1:10,000, Cell Signaling), anti-pIRF3 (1:500, 4D4G, Cell-Signaling), anti-IRF3 (1:2000, Cell-Signaling), anti-pTBK1 (1:500, D52C2, Cell-Signaling), anti-TBK1 (1:500, E9H5S, Cell-Signaling) overnight at 4 degrees Celsius and then washed with TBS-Tween (0.1%) the following day 3 times five minutes each wash. Secondary antibodies anti-mouse or rabbit-HRP (1:2000, Cell-Signaling) were probed for 1 hour at room temperature and then washed an additional three times for five minutes each wash. All images were processed and imaged on Bio-Rad’s chemidoc with chemiluminescence or by fluorescence.

Co-Immunoprecipitations experiments were lysed with pierce Co-IP lysis buffer (25 mM Tris-HCl pH 7.4, 150 mM NaCl, 1mM EDTA, 1% NP-40, 5% glycerol, Thermo-Fisher Scientific) containing pierce protease and phosphatase tablets inhibitor mini tablets, EDTA free (Thermo-Fisher scientific). 100ug of protein was incubated with anti-cGAS (1:50) overnight and the following day incubated with anti-Rabbit DynaBeads (Thermo-Fisher scientific) and then incubated 2 hours at four degrees Celsius. The following day samples were washed 5 times with Co-Ip lysis buffer and beads were boiled for five minutes in 2x Laemmli buffer and then separated on an 8% SDS-PAGE, transferred to a nitrocellulose membrane, and blocked with 5% BSA. Membranes were probed with anti-cGAS (1:1000) or anti-pp65 (1:200) antibody overnight at 4 degrees Celsius. The following day samples were probed with respective secondary, and membranes were developed in chemiluminescent buffer and imaged on a BioRad chemidoc.

### Biotin capture of nitrosylated proteins

100 ug of protein from HCMV-infected cell lysates was subjected to a biotin-switch assay following manufacturer protocol (Cayman Chemical Company). Biotinylated proteins were affinity purified with streptavidin-coated beads M-280 Dynabeads (Thermo-Fisher Scientific) overnight and washed 6 times with cold PBS using life technologies Dynamag-2. Beads were boiled in 2x leammli and loaded onto an 8% SDS-PAGE. Protein was separated and probed for antibodies specific to HCMV pp65.

### IFN-β1 ELISA

Cytokine secretion was measured using the “VeriKine Human IFN Beta ELISA Kit” from pbl assay science. Parental and pp65 expressing stable cell lines were transfected with 0.2 ug of G3-YSD for 8 hours in opti-mem. Media was collected, clarified and 50uL of supernatant was added to respective wells and incubated for 1 hour at room temperature. Wells were washed three times with ELISA wash buffer (TBS 0.05% Tween-20) and liquid aspirated from wells. Antibody from the kit was diluted 1:150 and antibody solution was added to wells and incubated for 1 hour at room temperature. Wells were then washed again as stated before and HRP-conjugate secondary antibody was diluted 1:100 and added to wells for 1 hour at room temperature. After another three washes, TMB substrate was added to the wells and allowed to develop for 15 minutes in the dark at room temperature. Once the ELISA plate fully developed stop solution was added to each well and absorbance measured at 450nm. Cytokine measurement was quantified by calculating pg/mL of IFN-β1 in the supernatant using the equation provided by the standard curve.

### Statistics

All experiments in this paper were analyzed using One-Way ANOVA and unpaired student T-test. Results were considered significant if the calculated p-Value was < 0.5 indicated by a 95% confidence interval. All data was graphed and analyzed utilizing the program Graph Pad Prism.

## Acknowledgments

We are grateful for Masatoshi Nukui in developing reagents for this project and initial work on discovering that HCMV proteins are protein-S-nitrosylated in HCMV infection. We thank Zsuzsa Szemere Ph.D., Meghan Carter, Emmanuel Ijeize Ph.D., and Mary Root for technical support and scientific advice.

